# Parabrachial nucleus activity in nociception and pain in awake mice

**DOI:** 10.1101/2023.03.22.533230

**Authors:** Jesse Andrew Smith, Yadong Ji, Rebecca Lorsung, Macauley Smith Breault, Jeffrey Koenig, Nathan Cramer, Radi Masri, Asaf Keller

## Abstract

The parabrachial nuclear complex (PBN) is a nexus for aversion, and for the sensory and affective components of pain perception. We have previously shown that, during chronic pain, PBN neurons in anesthetized rodents have amplified activity. We report a method to record from PBN neurons of behaving, head-restrained mice, while applying reproducible noxious stimuli. We find that both spontaneous and evoked activity are higher in awake animals, compared to urethane anesthetized mice. Fiber photometry of calcium responses from CGRP-expressing PBN neurons demonstrates that these neurons respond to nociceptive stimuli. In both males and females with neuropathic or inflammatory pain, responses of PBN neurons remain amplified for at least 5 weeks, in parallel with increased pain metrics. We also show that PBN neurons can be rapidly conditioned to respond to innocuous stimuli, after pairing with nociceptive stimuli. Finally, we demonstrate that changes in PBN neuronal activity are correlated with changes in arousal, measured as changes in pupil diameter.

**Significance Statement:** The parabrachial complex is a nexus of aversion, including pain. We report a method to record from parabrachial nucleus neurons of behaving mice, while applying reproducible noxious stimuli. This allowed, for the first time, tracking the activity of these neurons over time in animals with neuropathic or inflammatory pain. It also allowed us to show that the activity of these neurons correlates with arousal states, and that these neurons can be conditioned to respond to innocuous stimuli.

## Introduction

Pain is a multidimensional experience, composed of unpleasant sensory, affective and cognitive experiences (Melzack and Casey, 1968; Fields, 1999; Price, 2000; Treede et al., 2000; Auvray et al., 2010). Because of the multidimensionality of the pain experience, chronic pain patients suffer from comorbid emotional and cognitive deficits, and exhibit alterations in the function of brain networks linked to these deficits (Borsook, 2012; Bushnell et al., 2013). Therefore, treating pain, and especially chronic pain, requires addressing not only the sensory aspects of pain, but also its affective and cognitive components.

Chronic pain affecting the trigeminal system, in particular, is associated with deficits in affective processing, including increases in anxiety and depression (Smith et al., 2013; Wu et al., 2015; Turnes et al., 2022). A key region in the trigeminal system is the parabrachial nuclear complex (PBN). It is a nexus of aversion and of both the nociceptive and affective components of pain processing (Chiang et al., 2019). PBN is involved in other aversive behaviors (Cai et al., 2018), including taste aversion (Carter et al., 2015; Chen et al., 2018), threat memory (Han et al., 2015), and fear conditioning (Campos et al., 2018; Bowen et al., 2020). We and others have found that PBN neurons respond to noxious stimuli under urethane anesthesia (Bernard and Besson, 1990; Uddin et al., 2021) and display amplified activity during chronic pain (Uddin et al., 2018; Raver et al., 2020).

Most of what is known about the physiology of PBN in the context of pain comes from studies using anesthetized animals. This is problematic because anesthesia may blunt pain related activity. Further, anesthesia does not allow studies of state-dependent changes in pain perception, changes that are critical to pain behavior. Pain perception depends on both external stimuli and the internal state of the animal. Internal states—such as distraction (Buhle and Wager, 2010; Legrain et al., 2011) or hunger (Alhadeff et al., 2018)—profoundly alter pain perception. Therefore, it is essential to determine how internal states affect PBN processing in behaving animals.

Recordings of nociceptive responses from PBN in awake animals are complicated by at least two major technical difficulties. First, the location of PBN—at the junction of the midbrain and pons and its relation to major vessels—renders recordings from single units unstable and of low signal to noise ratio in behaving animals. The second complication is the ability to present to behaving animals reproducible noxious stimuli. We have adapted a head-restrained, awake rodent recording model to address these issues. This allowed us to investigate the physiology of PBN in awake animals in response to both noxious and non-noxious stimuli and the role of internal states in PBN processing.

## Methods

We adhered to accepted standards for rigorous study design and reporting to maximize the reproducibility and translational potential of our findings, as described by Landis et al. (2012) and in ARRIVE (Animal Research: Reporting In Vivo Experiments). In line with National Institutes of Health recommendations for scientific rigor, we performed an a priori power analysis to estimate required sample sizes (Landis et al., 2012).

### Animals

All procedures were conducted according to Animal Welfare Act regulations and Public Health Service guidelines and approved by the University of Maryland School of Medicine Animal Care and Use Committee. We used 27 male and female wild type C57BL/6 mice from Jackson Laboratory (JAX strain # 000644): 9 CCI-ION animals, 4 CFA injected animals, and 14 naïve animals. We also used 8 CGRP-CRE mice, 4 male and 4 female, that were bred in house from breeding pairs obtained from Jackson Laboratory (JAX stock # 033168).

Animals were housed in a 12 h dark/light cycle, and food and water were given ad libitum. The animals were group-housed prior to headplate surgery. After surgery they were individually housed.

### Headplate Implant Surgery

Animals were induced with 2-4% isoflurane and placed in a stereotaxic frame. The anesthesia was maintained at 1-2% for the surgery. Animals were given Rimadyl (5 mg/kg) for analgesia. The skin on the top of the skull was removed and the headplate (Model 11, Neurotar, Helsinki, Finland) was secured to the skull using a mixture of dental cement and Vetbond (3M, USA). Two screws were implanted anterior to the headplate for grounding. A craniotomy was created to expose the brain above PBN (AP 4.8mm-5.3mm, ML 0.5mm-2mm) and the dura was removed. The area was cleaned using a cortical buffer with antibiotics (125mM NaCl, 5mM KCl, 10mM Glucose, 10mM HEPES, 2mM 1M CaCl_2_, 2mM 1M MgCl_2_, 100 units/ml penicillin, 0.1 mg/ml streptomycin). The craniotomy was closed using a silicone elastomer (QuikSil, World Precision Instruments, Sarasota, FL, USA). The animal was placed on a heating pad and allowed to recover.

### Training

Animals were trained on the Neurotar Mobile HomeCage (Neurotar) where they were head restrained but allowed to locomote freely on a carbon fiber platform supported by a cushion of air. Animals were given 5 days to recover after the implant of the headplate. On day 6 animals were left in their home cage and placed near the Neurotar setup to habituate to the noise of the air flow for 2 hours. On day 7, animals were placed in the Neurotar setup and allowed to move freely for up to 2 hours to habituate to the device. Electrophysiology recordings started on day 8.

### Electrophysiology Recordings

#### Awake Electrophysiology Recordings

We performed recordings in low light conditions. Animals were placed in the Neurotar setup and allowed to acclimate for 5 minutes. The silicone elastomer was removed and the craniotomy was cleaned using a cortical buffer with antibiotics. A single platinum-iridium recording electrode (0.3-1 MΩ), produced in our lab, was lowered into PBN. We digitized recorded waveforms using a Plexon system (Plexon Inc., Dallas, TX). We isolated units that were responsive to noxious heat (50°C) applied to the anterior maxillary region of the head using a Picasso Lite dental surgical laser (AMD Lasers, Indianapolis, IN), positioned 5 mm above the skin, set to 3.5 W for 5 seconds. The laser was calibrated with a Jenco Electronics microcomputer thermometer to generate 51°C at the end of a 5 second exposure. We waited at least 2 minutes between consecutive stimuli and inspected the skin after each stimulus for erythema or tissue damage. Upon finding a neuron that was responsive to noxious heat stimulation, we allowed the cell to return to its baseline firing rate before recording baseline spontaneous activity for 5 minutes. After that, we recorded responses to noxious heat applied at 2 minute intervals to prevent tissue damage for a total of 10-12 trials. After each recording session, the electrode was removed and the craniotomy was cleaned using a cortical buffer with antibiotics and covered with fresh silicone elastomer.

#### Anesthetized Electrophysiology Recordings

Animals were anesthetized via intraperitoneal injections of urethane (10% w/v). Following anesthesia, the mice were placed in a stereotaxic frame with a heating pad and a craniotomy was made over the recording site to target PBN (AP 4.8mm-5.3mm, ML 0.5mm-2mm). The recording and stimulation protocol was identical to the awake mice protocol.

#### Electrophysiology Data Analysis

Recordings were sorted using Offline Sorter (Plexon Inc., Dallas, TX) using dual thresholds and principal component analysis. Responses to thermal stimuli were analyzed with custom MATLAB scripts. Significant responses were defined as firing activity exceeding the 95% confidence interval of the average pre-stimulus firing rate. Thus, evoked responses are expressed as significant responses above baseline firing levels. Peristimulus time histograms (PSTHs) were generated to analyze responses to repeated stimuli.

We defined after-discharges—periods of sustained activity that outlast a stimulus presentation (Okubo et al., 2013)—as PSTH bins in which activity exceeded the 95% confidence interval for a period lasting at least 5 seconds after stimulus offset.

### CCI-ION

We used a rodent model of neuropathic pain that was induced by chronic constriction of the infraorbital nerve (CCI-ION) (Bennett and Xie, 1988; Uddin et al., 2018; Raver et al., 2020). Animals were anesthetized with ketamine/xylazine (100/10 mg/kg, i.p.). We made intraoral incisions along the roof of the mouth, beginning distal to the first molar. The ION was freed from the surrounding connective tissue and loosely tied with a silk thread (4-0) 1-2 mm from the nerves emerging from the infraorbital foramen. Animals were monitored for 2 days in their home cage as they recovered. Each animal served as its own control, as recordings were compared before and after CCI in each animal.

### CFA

We used complete Freund’s adjuvant (CFA) to induce persistent inflammatory pain. Animals were induced with 2-4% isoflurane and injected with 15 µL of CFA (Sigma-Aldrich F5881. Sigma, St. Louis, MO, USA) subcutaneously in the snout, ipsilateral to the recording site. Animals were monitored for 24 hours in their home cage as they recovered before recording. Each animal served as its own control, as above.

### Behavior

#### Facial Grimace

To assess ongoing pain, we analyzed facial grimace behaviors in animals placed in the Neurotar Mobile HomeCage and images of the face were taken every 60 seconds for 15 minutes. Animals were scored with defined action units (AU) that examined orbital tightening, nose bulging, whisker positioning, and cheek bulging to derive a mouse grimace score (MGS) (Langford et al., 2010; Akintola et al., 2017). We calculated MGS as the average score of all the AUs.

#### Mechanical Sensitivity

To assess mechanical sensitivity, animals were placed in the Neurotar Mobile HomeCage and von Frey filaments (North Coast Medical) were applied to the anterior maxillary region, ipsilateral to CCI or CFA injections. A response was defined by an active swipe of the filament with the forepaws. We used the up-down method to determine response thresholds, as described previously (Dixon, 1965; Chaplan et al., 1994; Akintola et al., 2017; Raver et al., 2020). For each mouse, baseline response thresholds were determined prior to CCI-ION/CFA injections and thresholds were reassessed on each day of recordings from PBN. Hyperalgesia was defined as mechanical withdrawal threshold ≥ 20% below baseline.

### Viral Construct Injection

We anesthetized the animals with isoflurane and placed them in a stereotaxic frame. Either left or right PBN (-5.2mm AP, ±1.5mm ML, -2.2 to -2.9mm DV) was targeted via a small craniotomy. We injected 0.5 µL of AAV9-syn.Flex.GCaMP6f.WPRE. SV40 (Addgene #100833) at a rate of 0.1 µL/min using glass pipettes (40-60 µm tip diameter), coupled to a Hamilton syringe controlled by a motorized pump. The pipette was left in place for 10 minutes before being slowly retracted over 5-10 min. Headplates were implanted immediately after the virus injections.

Because AAV9 can be transported retrogradely (Castle et al., 2014), recorded signals might reflect activity in axons projecting to PBN. We carefully considered this possibility by analyzing histological sections through structures that project to PBN. We did not detect retrogradely labeled cells in any of these structures.

### Fiber Photometry

#### Recording

We anesthetized animals using 2% isoflurane and placed them in a stereotaxic frame. Mice were implanted with a fiber optic probe (400 µM diameter, 0.39NA; RWD Life Sciences) in the PBN (-5.2mm AP, +1.5mm ML, -2.2 to -2.5mm DV) during the same surgery they were injected with the viral construct and the head plate was implanted. The mice were given 3 weeks to recover and to allow for viral expression in their home cage.

For recordings, the fiber optic probe was connected to an RZX10 LUX fiber photometry processor running Synapse software (Tucker-Davis Technologies) through a Doric mini cube (Doric Lenses). LEDs at 465 nm and 405 nm were used for GCaMP excitation and isosbestic control, respectively. LED power was calibrated weekly using a digital optical power meter (Thor Labs).

#### Analysis

We used pMat (Bruno et al., 2021) to quantify photometry signals as changes in GCaMP fluorescence. These were converted to Z-Scores that were time-locked using a time window starting 5 seconds before the onset of the noxious stimulus and lasting until 30 seconds post-stimulus for each trial. A significant response was defined as one exceeding the 95% confidence interval of the baseline across all trials. The duration and AUC were calculated using the significant response of the Z-Score averaged across trials.

### Conditioning

#### Recordings

Mice were head restrained in the Neurotar Mobile Homecage and recordings were determined to be responsive to noxious heat (see above). While recording pupil size (see below) and firing rate of PBN neurons, we presented the mouse with a tone (1 second, 100 kHz, 88 dB) 10 times with 2 minutes intervals. We administered a conditioning paradigm where a 1 second tone (CS) was paired with a 5 second noxious heat stimulus (US), where the tone was presented 3 seconds after the onset of the noxious heat, to align with the laser reaching noxious temperature. This pairing was repeated 15 times at 1 minute intervals. The CS was presented until extinction with 2 minute between presentations of the CS. We then administered the conditioning paradigm a second time. The CS was again presented until extinction.

The firing rates of conditioned trials were binned at 0.1 seconds before being split into trials time-locked at the stimuli onset. A significant response was defined as exceeding the 95% confidence interval of the baseline across all trials.

#### Pupillometry

We videotaped pupils in head restrained mice, under low light conditions using a webcam (Logitech Brio, Logitech Inc., Newark, CA). Recordings were acquired at 30 Hz using custom MATLAB^®^ scripts and the Image Acquisition Toolbox (Mathworks, Natick, MA). Noxious heat stimulation timestamps were recorded by a microcontroller (Arduino Uno R3, 2015) using a custom script in Python (Python Software Foundation, https://www.python.org/). These timestamps aligned the video with the electrophysiological data. Pupil area was measured from the videos offline using Facemap software (Stringer et al., 2019). We excluded times when Facemap detected that the animal blinked. Pupil area was normalized between [0,1] using the minimum and maximum value within each recording session. Normalized pupil areas of conditioned trials were binned at 0.1 seconds before being split into trials time-locked at the stimuli onset. A significant response was defined as exceeding the 95% confidence interval of the baseline across all trials.

We defined spontaneous activity as activity at least 3 minutes before the onset of the first stimulus. Firing rates and normalized pupil areas during this time were binned using a width of 1 second. The normalized pupil areas were down-sampled to accommodate the bins of the spike rate by calculating the median within each bin. We applied a minimum cut-off of 10 Hz to the firing rates. To investigate the relationship between spontaneous activity and normalized pupil area we quantified it as the Pearson correlation coefficient.

To determine the relative time dynamics between firing rates and normalized pupil area after heat stimuli trials, we calculated a cross-correlation to produce lag values. This was done by binning the firing rates and normalized pupil areas using a width of 0.1 seconds and splitting them into trials, time-locked around the stimulus onsets. The lag that maximized the magnitude of correlation was selected for further analysis. The firing rates and normalized pupil area used for cross-correlation were then shifted using this lag before taking the Pearson correlation coefficient.

### Statistical analysis

We analyzed group data using GraphPad Prism version 9 for Mac (GraphPad Software, La Jolla CA) and custom scripts in MATLAB. Unless otherwise noted, data are presented as median ± 95% confidence intervals (95% CI). Unless otherwise indicated we used nonparametric Mann-Whitney *U* tests because data were not normally distributed. Individual statistical tests were run for each animal, as shown in the figures. We then averaged data from all animals for each condition to calculate group comparisons, and Cohen’s *d* was calculated for effect size. Correlations are Pearson correlations.

## Results

### Anesthesia suppresses spontaneous and evoked PBN responses

PBN neurons recorded in anesthetized animals respond robustly to nociceptive inputs (Bernard and Besson, 1990; Uddin et al., 2018; Raver et al., 2020; Uddin et al., 2021). We developed an approach to apply reproducible noxious stimuli to awake, head-restrained mice, while recording well-isolated single units from the PBN nuclear complex (PBN; Figs. 1, 2). Nociceptive stimuli were generated by a laser-generated light spot (3mm diameter) applied to the snout. This generated a heat stimulus that increased—during the 5 second stimulus period—from body temperature to 50°C (red bar in Fig. 1A, B). Figure 1 compares examples of peri-event histograms recorded from individual PBN neurons in an anesthetized mouse (Fig. 1A) and in an awake mouse (Fig. 1B). In the neuron from the awake animal, spontaneous activity, shown before the onset of the noxious heat, is markedly higher than that in the anesthetized animals. The amplitude of the response is also larger in the neuron from the awake animal.

**Figure 1:**
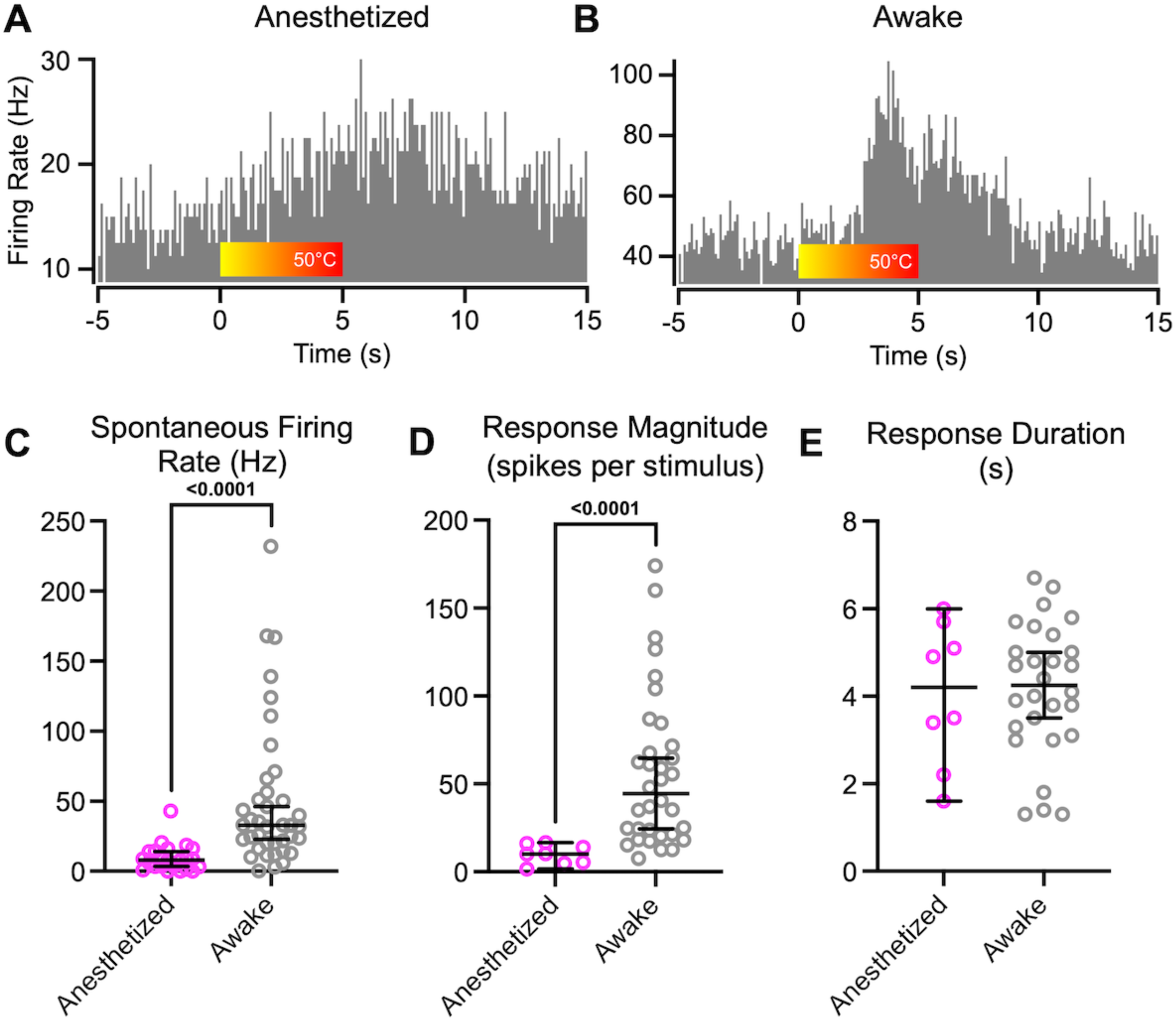
PBN neurons in anesthetized mice respond substantially differently than in awake animals. A: PSTHs recorded from PBN neurons in a urethane-anesthetized and awake (B) mice. Color bar indicates application of heat from a laser, producing temperatures that peak at 50°C. Note different y-axes scales. C: Spontaneous firing rates and magnitude of responses to noxious heat applied to the face (D) were higher in awake mice (Mann-Whitney U = 109.5, p < 10-4 and U = 13, p < 10-4). E: Response durations were indistinguishable in awake and anesthetized mice. Data are medians and 95% confidence intervals.

**Figure 2:**
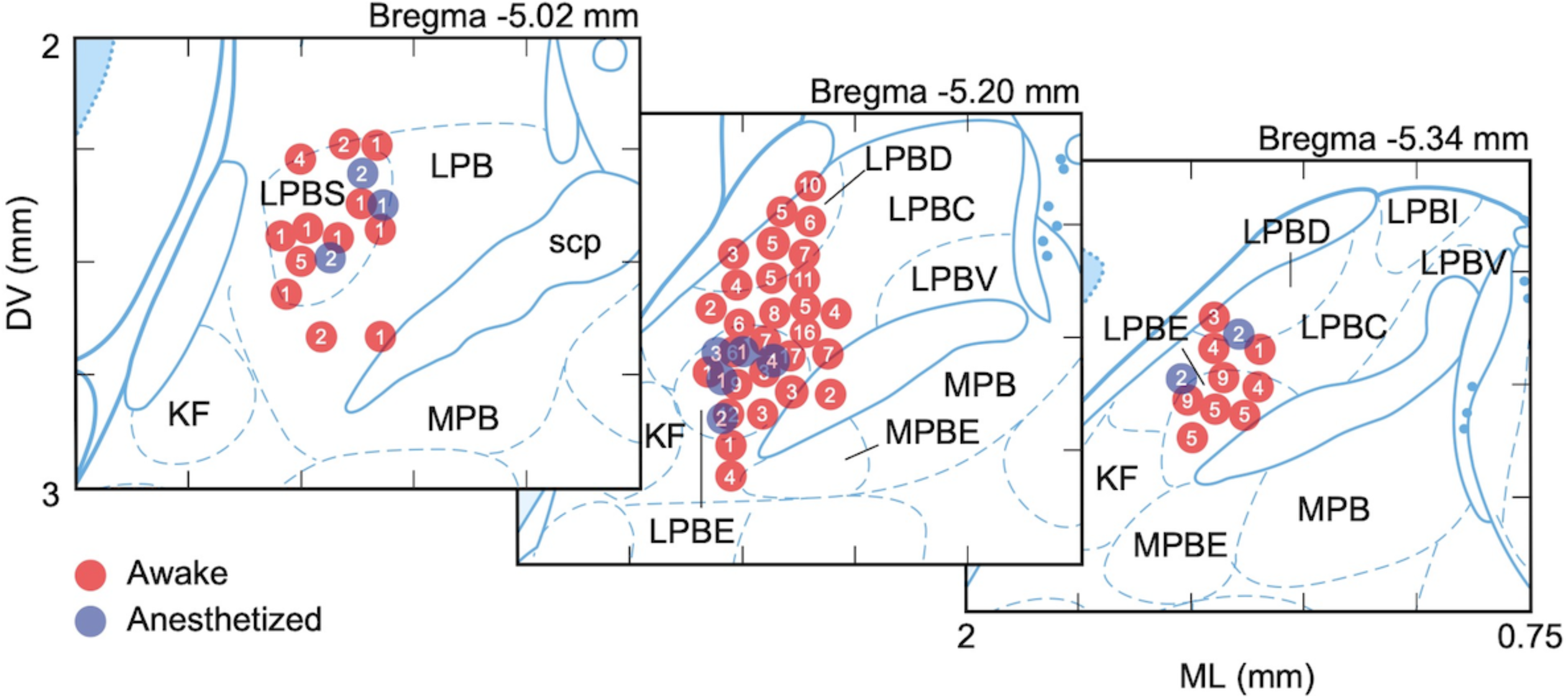
Recordings sites in PBN. Locations of recording sites, derived from stereotaxic coordinates, indicated by red markers (for neurons recorded from awake animals) or blue markers (for neurons recorded from anesthetized animals). Numbers indicate the number of recordings from the same location. Kölliker-fuse nucleus (KF), lateral PBN (LPB), dorsal part of the lateral PBN nucleus (LPBD), central part of the lateral PBN nucleus (LPBC), external part of the lateral PBN nucleus (LPBE), internal part of the lateral PBN nucleus (LPBI), superior part of the lateral PBN nucleus (LPBS), ventral part of the lateral PBN nucleus (LPBV), medial PBN nucleus (MPB), external part of the medial PBN nucleus (MPBE), superior cerebellar peduncle (scp). Image maps from Paxinos & Franklin, 2019.

Group data comparisons quantify the differences in response properties in anesthetized and awake mice (Fig. 1C-E). Spontaneous firing rates were, on average, more than 5 times higher in awake mice (Fig. 1C, Cohen’s d 1.09, p < 10^-4^, Table 1). As detailed in Methods, we considered significant responses spiking activity that exceeded the 95^th^ percent confidence interval of the average pre-stimulus firing rate. We computed response magnitude and duration of these significant response epochs. Response magnitudes (computed as spikes per stimulus per bin) were, on average more than 5 times higher in awake mice (Fig. 1D, Cohen’s d 1.49, p = 0.003, Table 1), compared to those in anesthetized mice. Response durations were similar in neurons from awake and anesthetized mice (Fig. 1E, p > 0.99, Table 1).

**Table 1:** Statistics for Figures 1 through 9 showing the corresponding figure number, animals, metric, comparisons being made, test statistic, medians or means, sample size, and *p* values.

| Figure | Animal | Metric | Comparison | Test Statistic | Median | Sample Size | P value |
| --- | --- | --- | --- | --- | --- | --- | --- |
| 1C | N/A | Spontaneous activity | Anesthetized vs awake | Mann-Whitney<br>U = 109.5 | Anesthetized= 7.94<br>Awake= 32.80 | Anesthetized= 23 cells<br>Awake= 39 cells | <10<sup>-4</sup> |
| 1D | N/A | Response Magnitude | Anesthetized vs awake | Mann-Whitney<br>U = 13 | Anesthetized= 10.15<br>Awake= 44.35 | Anesthetized= 23 cells<br>Awake= 28 cells | <10<sup>-4</sup> |
| 1E | N/A | Response Duration | Anesthetized vs awake | Mann-Whitney<br>U = 112 | Anesthetized= 4.20<br>Awake= 4.25 | Anesthetized= 8 cells<br>Awake= 28 cells | >0.99 |
| 4E | F6 | Spontaneous activity | Pre vs Post CCI | Mann-Whitney<br>U = 18 | Pre= 20.50<br>Post= 32.80 | Pre= 8 cells<br>Post= 7 cells | 0.28 |
|  | M9 |  |  | Mann-Whitney<br>U = 14 | Pre= 7.47<br>Post= 23.25 | Pre= 11 cells<br>Post= 6 cells | 0.058 |
|  | F7 |  |  | Mann-Whitney<br>U = 41 | Pre= 35.35<br>Post= 25.15 | Pre= 8 cells<br>Post= 12 cells | 0.624 |
|  | M12 |  |  | Mann-Whitney<br>U = 56 | Pre= 43.90<br>Post= 35.00 | Pre= 8 cells<br>Post= 12 cells | 0.691 |
| 4F | F6 | Response<br>Duration | Pre vs Post CCI | Mann-Whitney<br>U = 3.5 | Pre= 3.20<br>Post= 9.40 | Pre= 6 cells<br>Post= 7 cells | 0.009 |
|  | M9 |  |  | Mann-Whitney<br>U = 0 | Pre= 4.25<br>Post= 9.50 | Pre= 6 cells<br>Post= 5 cells | 0.0016 |
|  | F7 |  |  | Mann-Whitney<br>U = 3 | Pre= 4.40<br>Post= 11.90 | Pre= 8 cells<br>Post= 11 cells | 0.0002 |
|  | M12 |  |  | Mann-Whitney<br>U = 0 | Pre= 4.70<br>Post= 10.70 | Pre= 7 cells<br>Post= 18 cells | <10<sup>-4</sup> |
| 4G | F6 | Response<br>Magnitude | Pre vs Post CCI | Mann-Whitney<br>U = 4 | Pre= 24.62<br>Post= 50.35 | Pre= 6 cells<br>Post= 7 cells | 0.1143 |
|  | M9 |  |  | Mann-Whitney<br>U = 12 | Pre= 23.70<br>Post= 98.65 | Pre= 6 cells<br>Post= 5 cells | 0.0353 |
|  | F7 |  |  | Mann-Whitney<br>U = 17 | Pre= 58.90<br>Post=307.00 | Pre= 8 cells<br>Post= 11 cells | 0.0125 |
|  | M12 |  |  | Mann-Whitney<br>U = 2 | Pre= 64.60<br>Post= 374 | Pre= 7 cells<br>Post= 18 cells | <10<sup>-4</sup> |
| 6C | N/A | Spontaneous activity | Anesthetized vs awake Post CCI | Mann-Whitney<br>U = 69.50 | Anesthetized= 11.76<br>Awake= 32.80 | Anesthetized= 10 cells<br>Awake= 39 cells | 0.0012 |
| 6D | N/A | Response Duration | Anesthetized vs awake Post CCI | Mann-Whitney<br>U = 115.5 | Anesthetized= 16.70<br>Awake= 11.15 | Anesthetized= 10 cells<br>Awake= 38 cells | 0.0586 |
| 6E | N/A | Response Magnitude | Anesthetized vs awake Post CCI | Mann-Whitney<br>U = 85 | Anesthetized= 65.70<br>Awake= 282.00 | Anesthetized= 10 cells<br>Awake= 41 cells | 0.0034 |
| 7D | M10 | Spontaneous activity | Pre vs Post CFA | Mann-Whitney<br>U = 1 | Pre= 2.95<br>Post= 46.70 | Pre= 5 cells<br>Post= 7 cells | 0.0051 |
|  | F14 |  |  | Mann-Whitney<br>U = 25 | Pre= 29.80<br>Post= 70.20 | Pre= 15 cells<br>Post= 12 cells | 0.0009 |
|  | F15 |  |  | Mann-Whitney<br>U = 34 | Pre= 28.05<br>Post= 46.70 | Pre= 14 cells<br>Post= 10 cells | 0.0358 |
|  | M16 |  |  | Mann-Whitney<br>U = 14 | Pre= 26.00<br>Post= 52.00 | Pre= 8 cells<br>Post= 13 cells | 0.0045 |
| 7E | M10 | Response<br>Duration | Pre vs Post CFA | Mann-Whitney<br>U = 0 | Pre= 3.50<br>Post= 12.50 | Pre= 5 cells<br>Post= 7 cells | 0.0043 |
|  | F14 |  |  | Mann-Whitney<br>U = 3 | Pre= 3.90<br>Post= 9.10 | Pre= 16 cells<br>Post= 12 cells | <10<sup>-4</sup> |
|  | F15 |  |  | Mann-Whitney<br>U = 2 | Pre= 2.10<br>Post=9.18 | Pre= 15 cells<br>Post= 10 cells | <10<sup>-4</sup> |
|  | M16 |  |  | Mann-Whitney<br>U = 0 | Pre= 3.78<br>Post= 9.60 | Pre= 8 cells<br>Post= 13 cells | <10<sup>-4</sup> |
| 7F | M10 | Response<br>Magnitude | Pre vs Post CFA | Mann-Whitney<br>U = 0 | Pre= 11.92<br>Post=136.00 | Pre= 5 cells<br>Post= 7 cells | 0.0025 |
|  | F14 |  |  | Mann-Whitney<br>U = 2 | Pre= 17.20<br>Post=354.30 | Pre= 13 cells<br>Post= 12 cells | <10<sup>-4</sup> |
| | F15 | | | Mann-Whitney<br>U = 0 | Pre= 36.60<br>Post=317.50 | Pre= 11 cells<br>Post= 8 cells | $<10^{-4}$ |
|  | M16 |  |  | Mann-Whitney<br>U = 1 | Pre= 67.60<br>Post=701.50 | Pre= 7 cells<br>Post= 10 cells | 0.0002 |
| 8B | N/A | N/A | Firing rate vs Pupil<br>size correlation | Spearman's r<br>=0.542 | N/A | N/A | $2.36e^{-24}$ |

### PB^CGRP^ neurons are activated in response to acute pain

PBN neurons that respond to nociceptive stimuli, and whose activity is related to chronic pain conditions, are in the lateral portion of the PBN (Uddin et al., 2018; Raver et al., 2020). Many neurons in this region express calcitonin gene-related peptide (CGRP), a peptide implicated in aversive behaviors, including pain (Campos et al., 2018; Palmiter, 2018; Chiang et al., 2020). To test if PBN neurons that express CGRP (PB^CGRP^ neurons) respond to noxious stimuli, we used in vivo calcium imaging with fiber photometry to record population activity of PB^CGRP^ neurons.

Figures 3A-C depict GCaMP expression in PB^CGRP^ neurons of a male (B) and female (C) mouse, and the location of the optical fiber (A). An example of a heat-evoked CGaMP response is shown in Figure 3D. The portions of the response that exceeds the 95% confidence interval above the mean baseline levels, indicated by the dotted horizontal line, was defined as significant and labeled in green. The color bar shows the increase in heat, from room temperature to 50°C, which is just above the heat pain threshold for these animals. This threshold was determined based on published data, and on the known activation threshold of relevant nociceptors (Yeomans and Proudfit, 1996; Deuis et al., 2017).

**Figure 3:**
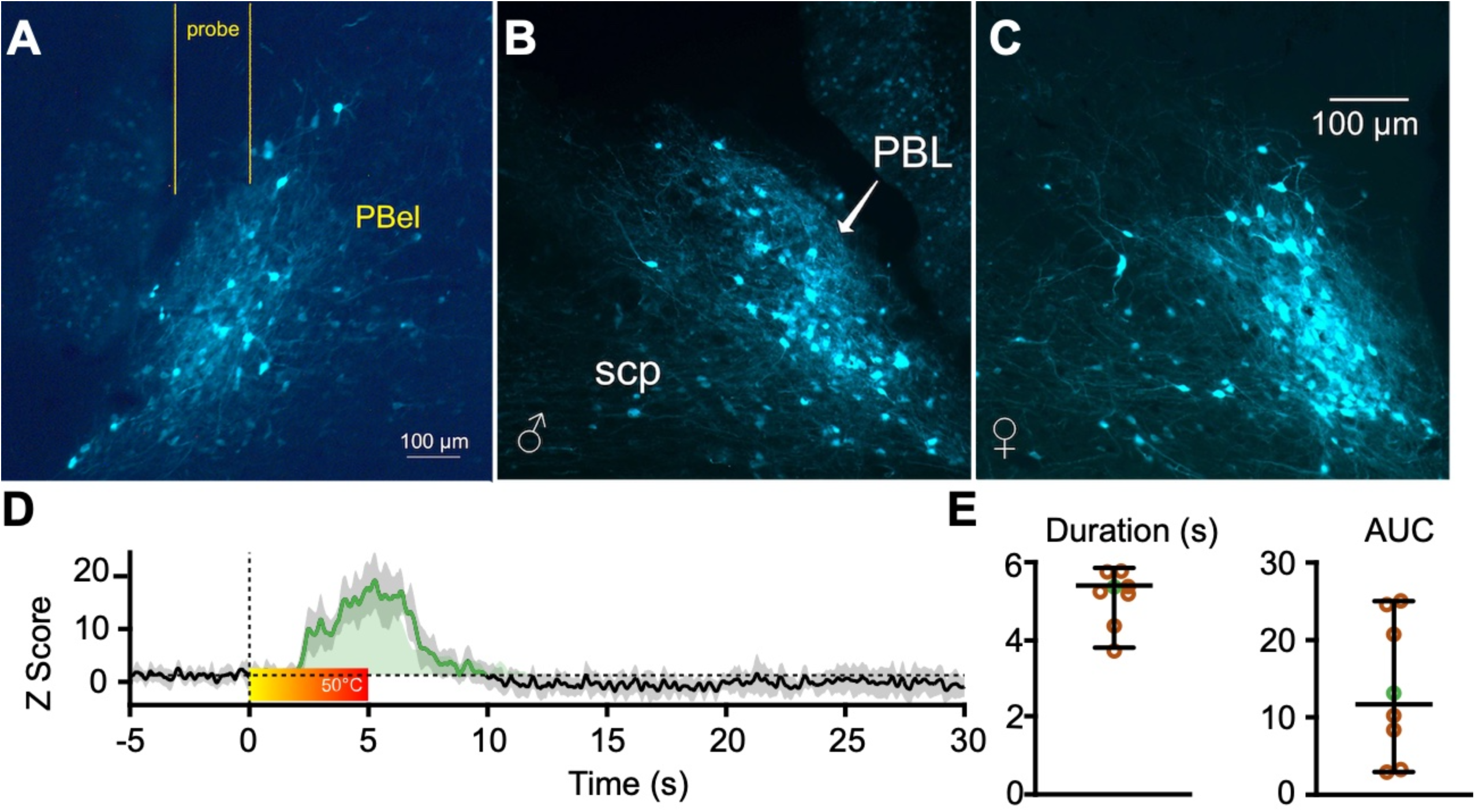
PB^CGRP^ neurons respond to noxious stimuli. GCaMP expression in PB^CGRP^ neurons of a male (B) and female (C) mouse, and the location of the optical fiber (A). D: Heat-evoked GCaMP response, with significant portion of the response—exceeding the 95% confidence interval above the baseline levels—indicated by dotted horizontal line and labeled in green. Color bar shows the increase in heat, from room temperature to 50°C. E: Response duration and area under the curve (AUC) of these responses, from 8 animals (4 of each sex). Data from the animal shown in D are marked in green circles.

We averaged responses recorded over 4 days from each animal. The duration and area under the curve (AUC) of these responses, from 8 animals (4 of each sex), are depicted in Figure 3E (data from the animal shown in Fig. 3D are marked in green circles in Fig. 3E). This experiment was not powered to test for sex differences. These findings demonstrate that PB^CGRP^ neurons respond robustly to noxious heat stimuli applied to the trigeminal region.

### Neuropathic pain is associated with hyperactivity of PBN neurons in awake animals

We, and others, have shown—in anesthetized rodents—that the activity of PBN neurons is amplified in models of chronic pain (Matsumoto et al., 1996; Uddin et al., 2018; Raver et al., 2020). We tested whether similar changes occur also in awake animals. As in our previous studies (Uddin et al., 2018; Raver et al., 2020), we used the chronic constriction injury of the infraorbital nerve (CCI) model, developed by Bennet and Xie (1988), to induce persistent pain-like behaviors. We tested mechanical hypersensitivity of the face in head restrained mice using a modified version of the “up-down method” (Chaplan et al., 1994) (see Methods). In each of the four animals with CCI, mechanical withdrawal thresholds were lower than in pre-CCI levels (Fig. 4A), indicating that the mice experienced increased tactile hypersensitivity. These changes were recorded between 8 and 56 days after CCI surgery.

**Figure 4:**
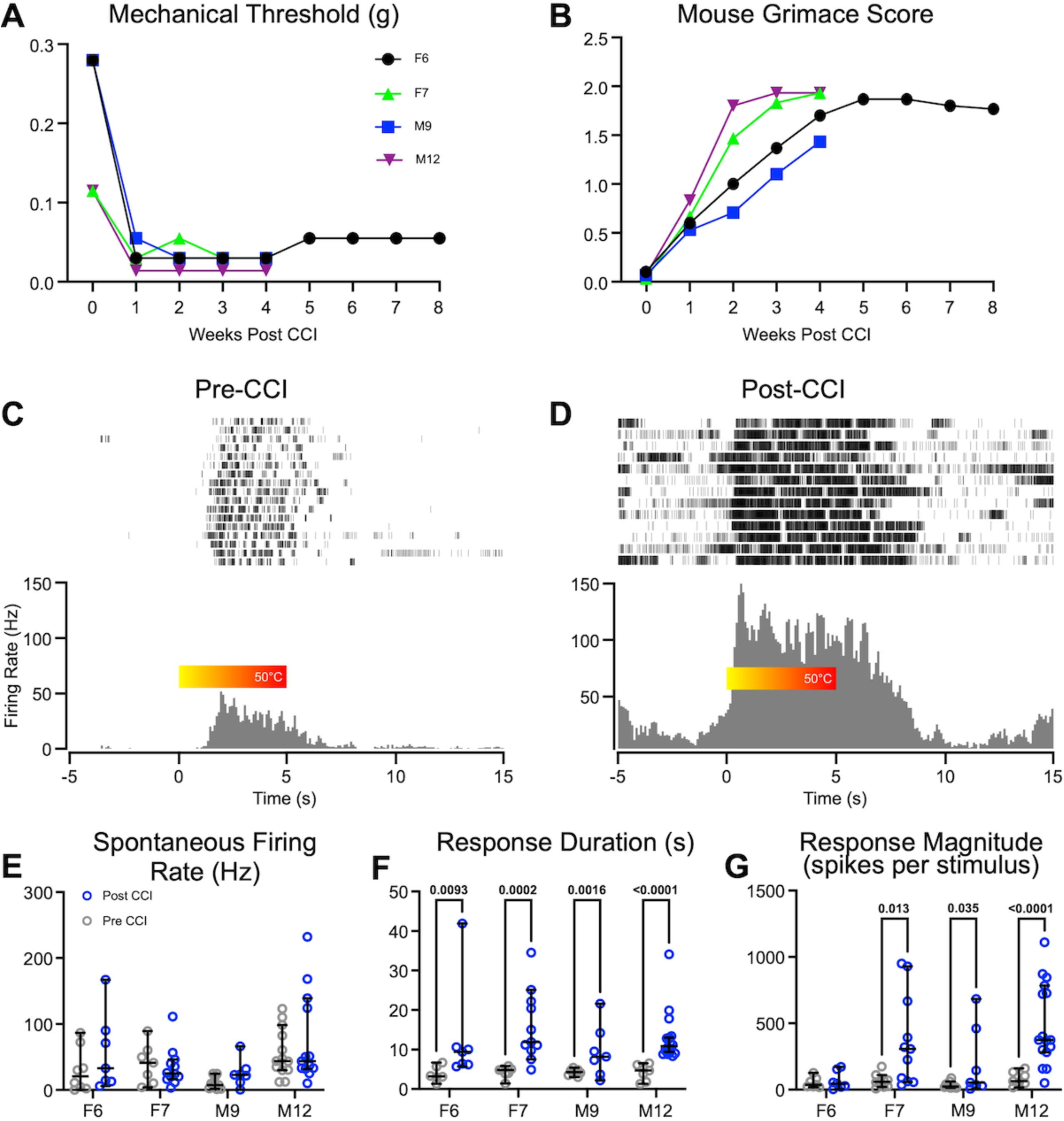
Neuropathic pain is associated with hyperactivity of PBN neurons in awake animals. Time course of reductions in facial tactile hypersensitivity (A) and increases in mouse grimace scores (B) in each of the four mice (M=male; F=female), after CCI. Representative raster plots and PSTHs of neurons recorded from the same animal before (C) and after (D) CCI, in response to noxious heat applied to the snout. Color bar shows the increase in heat from room temperature to 50°C. In each of the mice spontaneous activity was not affected after CCI (E), whereas response durations (F) and magnitudes (G) increased. Data represent medians and 95% confidence intervals; p-values derived from Mann-Whitney tests.

We used mechanical hypersensitivity as a behavioral readout because the thermal hypersensitivity tests of the face were unreliable, as even 0.1 W power differences in the calibrated laser output—the finest differences in power output the laser could produce—resulted in fluctuating behavioral responses. This may be due to the high sensitivity of the face to the thermal stimulation. Mechanical sensitivity tests, on the other hand, produced consistent behavioral responses at each filament level.

To assess ongoing pain, we used the mouse grimace scale (MGS) (Langford et al., 2010; Akintola et al., 2017). MGS scores were higher in mice after CCI, relative to controls, and remained higher for as long as they were recorded (up to 8 weeks) (Fig. 4B), suggesting that CCI mice were exhibiting persistent, ongoing pain.

We compared recordings of well isolated, single units from PBN in the same mice before and after CCI. There were distinct changes in the response magnitude and duration, but not the spontaneous activity, of PBN neurons after CCI (Fig. 4 E-G). Figure 4 shows an example of peri-event rasters and peri-event histograms recorded from individual PBN neurons in the same animal before (Fig. 4C) and 7 days after CCI (Fig. 4D). The neuron recorded after CCI showed an increase in both the magnitude and the duration of responses to heat stimuli applied to the face. In this neuron, spontaneous activity is also higher after CCI, however, when averaged across animals, spontaneous activity did not change after CCI (see below). The neurons recorded before CCI began responding at approximately 45°C, presumably the nociceptive threshold. In contrast, the neuron recorded after CCI began to fire at temperatures well below noxious threshold, at approximately 30°C (Fig. 4C and D). Before CCI, all neurons from all animals responded only to nociceptive temperatures; after CCI, all neurons also responded to innocuous temperatures.

We quantified the differences in response properties before and after CCI for each of the mice (Fig. 4E-G). We included, as post-CCI data, neurons recorded from two to eight weeks after CCI, a period in which all animals displayed tactile hypersensitivity and increased MGS scores. There was no difference in spontaneous firing rates after CCI in any of the animals (Fig. 4E), but there was an increase in the magnitude of the response (Fig. 4G, Table 1) and response duration (Fig. 4F, Table 1). The magnitude of the responses was 3 times larger after CCI (Fig. 4G, effect size 0.5, p < 10^-4^) and response durations were 3.25 times longer (Fig. 4F, effect size 0.5, p < 10^-4^). Median response duration after CCI was 10.5 seconds, far outlasting the duration of the stimulus. These prolonged responses are defined as after-discharges (Matsumoto et al., 1996; Uddin et al., 2018; Raver et al., 2020). The increase in response duration is consistent with our previous findings, in anesthetized rodents, demonstrating that chronic pain is associated with an increase in the incidence and duration of after-discharges in PBN neurons (Uddin et al., 2018; Raver et al., 2020).

We investigated sex differences by pooling 90 neurons recorded from 8 animals (4 of each sex). In naïve animals, there were no sex differences in spontaneous activity (p = 0.4, median (males)= 24.65, CI (males) = 20.51,42.72; median (females) = 29.95, CI (females) = 24.38,38.66), response duration (p = 0.84, median (males) = 4.8, CI (males) = 7.767, 20.94; median (females) = 5.3, CI (females) = 7.971, 17.63) or response magnitude (p = 0.9, median (males) = 37.95, CI (males) = 53.29, 166.6; median (females) = 35.05, CI (females) = 41.97, 76.39). Similarly, there were no sex differences after CCI in spontaneous activity (p = 0.34, median (males) = 32.45, CI (males) = 28.57, 77.52; median (females) = 25.9, CI (females) = 22.15, 62.45), response duration (p = 0.97, median (males) = 10.2, CI (males) = 10.09, 15.25; median (females) = 10.85, CI (females) = 9.53, 19.89), or response magnitude (p = 0.07, median (males) = 371.5, CI (males) = 292, 564.8; median (females) = 154, CI (females) = 108.5, 422.4).

### Time-course of changes in PBN activity

By recording from awake animals, we were able to investigate the time course of the activity of PBN neurons before and after the induction of nerve injury, and to determine the relationship between the activity of PBN neurons and the pain behaviors in individual animals. Figure 5 plots the duration (Fig. 5A) and magnitude (Fig. 5B) of responses in each of the neurons, pooled from the 4 animals, across time. Response duration and magnitude remained elevated relative to pre-CCI levels for at least 5 weeks post-CCI period, a period in which the animals experienced hyperalgesia (Fig. 5A). We recorded a small number of neurons from animals 6 to 8 weeks post-CCI; although some of these neurons had elevated activity (Fig. 5A, B), the small sample size precludes conclusions about activity levels at these time points.

**Figure 5:**
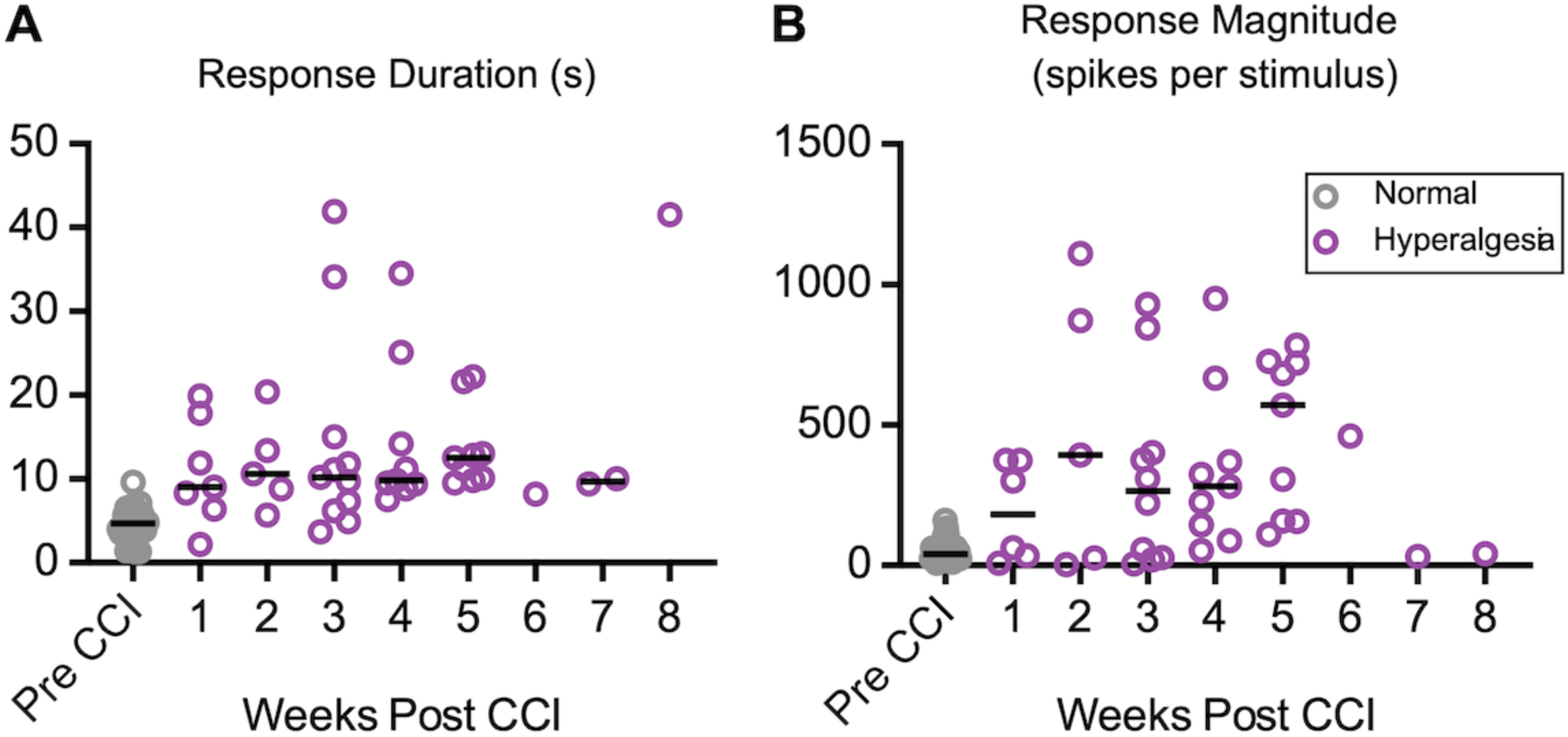
Time course of changes in PBN activity after CCI. Response durations (A) and magnitudes (B) remain elevated for at least 5 weeks post CCI (B). Purple markers indicate neurons recorded while animals displayed tactile hyperalgesia. Individual data represent individual neurons, and black bars depict weekly median values.

### Anesthesia suppresses the effects of CCI on PBN activity

We showed, above, that anesthesia suppresses the responses of PBN neurons to acute nociceptive inputs (Fig. 1). We also showed that CCI results in amplification of PBN responses, recorded from awake rodents (Fig. 4). Here, we tested if this amplification was affected by anesthesia. That is, we directly compared PBN activity after CCI, in anesthetized vs awake mice.

Group data comparisons quantified the differences in responses to noxious heat stimuli in anesthetized and awake mice after CCI (Fig. 6). Spontaneous firing rates were more than 3 times higher in awake mice (Fig. 6A, effect size 4.5, p = 0.03). Response magnitudes were more than 5 times higher in awake mice (Fig. 6C, effect size 5.5, p = 0.003). Response durations were similar in neurons from awake and anesthetized mice (Fig. 6B; p > 0.99). These findings indicate that spontaneous activity and responses to nociceptive stimuli in PBN neurons of CCI mice are larger in the awake, compared to anesthetized animals, but that the duration of after-discharges is similar in these conditions.

**Figure 6:**
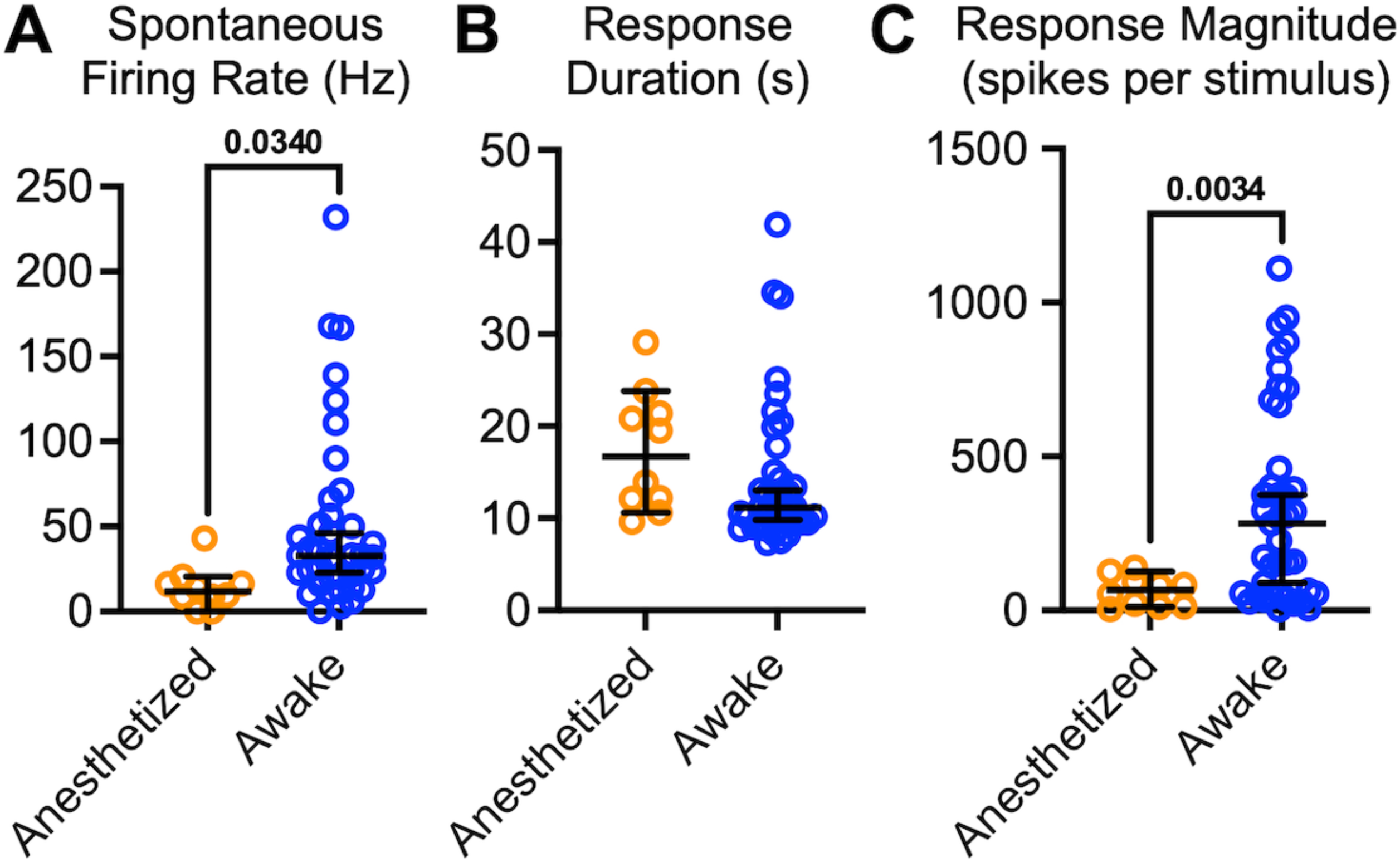
PBN neurons after CCI are different in awake animals. A: Post-CCI spontaneous firing rates recorded from awake mice were higher than in anesthetized mice. B: There was no difference in response duration in response to noxious heat applied to the face. C: Response magnitude was higher in awake mice than in anesthetized mice in response to noxious heat applied to the face. Markers represent cells (n = 10 cells for anesthetized and 44 for awake). Data represent medians and 95% confidence intervals.

### Inflammatory pain is associated with hyperactivity of PBN neurons in awake animals

To determine if similar changes in PBN activity occur also in other models of pain, we induced, in a separate group of animals, inflammatory pain by injection of complete Freund’s adjuvant (CFA) in the snout. We assessed tactile hypersensitivity levels at the snout daily on each of the 3 days before CFA injections, and on days 2 to 6 after injections. After CFA, mechanical thresholds decreased (Fig. 7A) indicating that the mice were experiencing increased tactile hypersensitivity.

**Figure 7:**
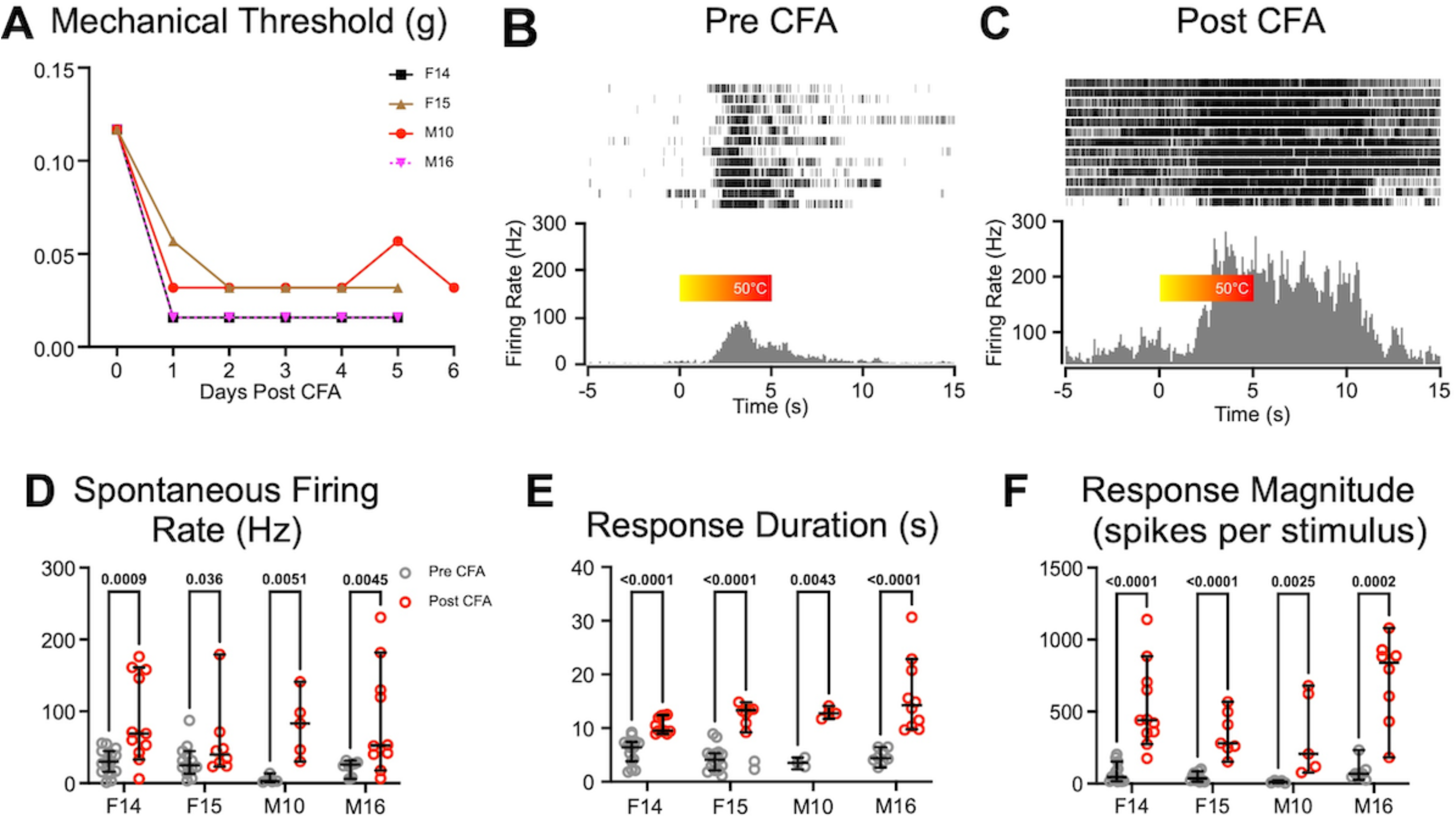
Inflammatory pain is associated with hyperactivity of PBN neurons in awake animals. A: Time course of reductions in facial tactile hypersensitivity in each of the four mice (M=male; F=female), after CFA injection. Representative rasters and PSTHs computed for neurons recorded from the same animal before (B) and after (C) CFA injection, in response to noxious heat applied to the snout for 5 seconds. The color bar shows the increase in heat from room temperature to 50°C. D: Spontaneous firing rates increased in 3 out of the 4 animals after CFA injections. Both response duration (E) and magnitude (F) increased after CFA in each of the animals. Data represent medians and 95% confidence intervals.

Between 2 and 6 days after CFA application, there were changes in the spontaneous and evoked activity of PBN neurons. Figure 7 depicts an example of peri-event rasters and peri-event histograms recorded from individual PBN neurons in the same animal before (Fig. 7B) and after CFA (Fig. 7C). After CFA, there were increases in spontaneous activity, response duration, and response magnitude in all animals tested (Fig.7D-F).

We quantified the differences in response properties before and 6 days after CFA in each of the awake mice (Fig. 7D-F). Unlike in CCI animals, there was an increase in spontaneous firing rates after CFA (Table 1). Neurons from all animals were pooled into pre and post CFA groups, revealing an average 3.3-fold increase in spontaneous activity (Fig. 7D, effect size 0.49, p < 10^-4^). The increase in spontaneous activity may reflect the persistent inflammatory nature of CFA pain, which may not be as prominent in animals with CCI. Just as with the CCI animals, response durations were 2.7 times longer (Fig. 7E, effect size 0.49, p < 10^-4^) and response magnitudes were 2.7 times larger after CFA (Fig. 7F, effect size 0.52, p < 10^-4^) in all animals tested, indicating the presence of amplified after discharges. Taken together with the CCI data, these data indicate that mouse models of both neuropathic pain (CCI) and inflammatory pain (CFA) are associated with hyperactivity of PBN neurons in awake mice.

We investigated sex differences by pooling 42 neurons recorded all 4 animals (2 males and 2 females). In animals recorded after CFA injections, there were no differences in spontaneous activity (p = 0.59, median (males) = 51.7, CI (males) = 43.94, 100.7; median (females) = 61.9, CI (females) = 51.64, 98.86), response duration (p = 0.13, median (males) = 12.3, CI (males) = 10.54, 16.21; median (females) = 10.7, CI (females) = 10.04, 11.94), or response magnitude (p = 0.92, median (males) = 432.4, CI (males) = 307.8, 662.3; median (females) = 340, 566.9).

### PBN activity is correlated with arousal

A benefit of recording from awake animals is the ability to test the hypothesis that PBN activity can be modulated by the “behavioral state” of an animal. One approach to assess behavioral states is to measure pupil size as a proxy of the arousal state (de Gee et al., 2020).

Spontaneous activity rates were correlated with pupil area. Figure 8A depicts a representative recording session, plotting changes in both pupil area (blue) and the spontaneous firing rate of a PBN neuron (red). Note that pupil dilations appear to immediately precede increases in the neuronal firing rate. Figure 8B quantifies this relationship by plotting instantaneous neuronal firing rates (1 second bins) against pupil are, demonstrating a positive correlation between these metrics. A significant, positive correlation was seen in 14 out of 17 cells across all 7 animals tested (4 males, 3 females), with r values ranging from 0.55 to 0.12. These findings suggest that arousal states—approximated by changes in pupil diameter—correlate with changes in PBN activity.

**Figure 8:**
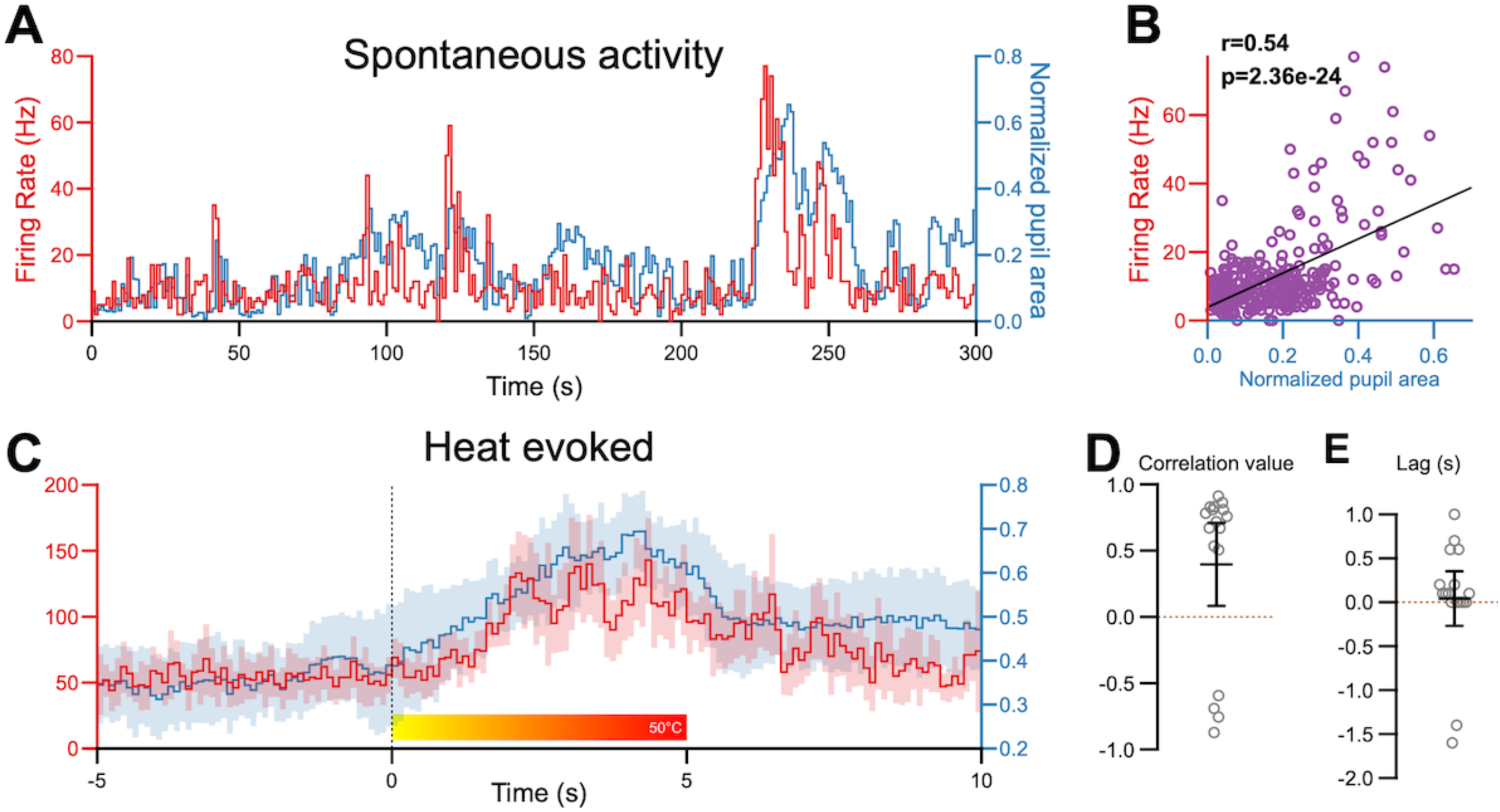
PBN activity is correlated with arousal states. A: Representative traces depicting spontaneous firing rates (red) of PBN neurons and normalized pupil area (blue). B: These metrics were positively correlated. C: Averaged (and 95% CI) traces of firing rate of PBN neurons (red) and pupil area (blue) in response to noxious heat applied to the snout. D: The lag-shifted correlation values suggest that firing rate and pupil area changes are positively correlated (n = 17 cells, 7 animals). E: The positive lag value after cross-correlation suggests that changes in firing rate preceded changes in pupil area (n = 17 cells, 7 animals).

To determine if arousal states affect evoked responses of PBN neurons, we constructed and overlayed PSTHs (0.1 second bin size) of neuronal activity and pupil area (Fig. 8C). The PSTHs were aligned to the onset of the 5 second heat stimuli applied to the snout. Pupil dilation and PBN activity after the onset of the noxious heat were highly correlated (Fig. 8C,D, r = 0.83, p = 1.75 x 10 ^-13^). A significant positive correlation was seen in 13 neurons from 7 animals tested, with r values ranging from 0.9 to 0.5. Four cells had a negative correlation (Fig. 8D). Cross correlation analysis revealed that, in the majority of cells recorded (15 of 17) there was a positive lag, indicating that changes in PBN activity preceded changes in pupil area (Fig. 8E).

### Auditory conditioning of PBN activity

Pairing activation of PBN neurons with sucrose produces robust and long-lasting conditioned taste aversion in mice, whereas suppressing PBN activity suppresses the acquisition and expression of this conditioned aversion (Carter et al., 2015; Chen et al., 2018). These findings suggest that the activity of PBN neurons can be conditioned to respond to innocuous stimuli. To test this, we conditioned head-restrained mice by pairing an auditory tone (1 second, 100 kHz, 80 dB) with a noxious heat stimulus applied to the snout. We did this while recording from nociceptive neurons in the PBN nucleus. Figure 9 depicts an example of one of these recording sessions.

**Figure 9:**
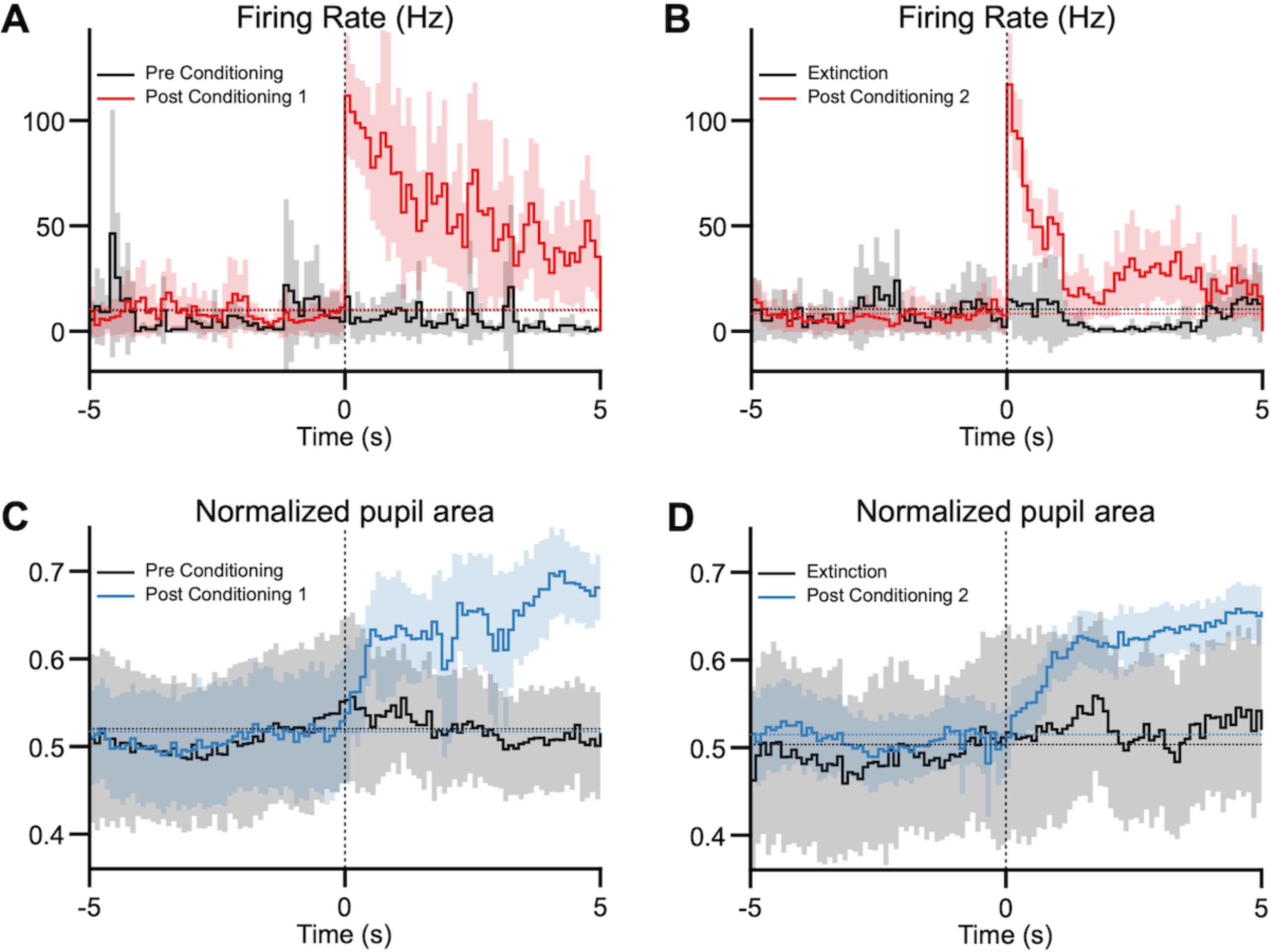
Innocuous auditory conditioning of PBN responses. A: A noxious heat-responsive PBN neuron does not respond to an innocuous tone before conditioning (black trace) but responds robustly to the tone after a conditioning (red). B: Representative traces showing the same PBN neuron undergoing extinction (black), and responses after a second conditioning paradigm (red). C: Similarly, pupil responses to the innocuous auditory stimulus appear only after the conditioning paradigm (blue). D: Recordings from the same neuron after extinction of the conditioned responses (black trace) and after reconditioning the auditory response (blue). Data represent averages and 95% confidence intervals, indicated by the horizontal dotted lines.

Before conditioning, the PBN neuron responded to the heat stimulus (not shown), but not to the auditory stimulus (Fig. 9A). We used 15 pairings of auditory tone (1 second) and heat (5 seconds) stimuli, with the auditory stimulus presented 3 seconds after the onset of the noxious heat stimulus, so that the auditory stimulus is presented just before the heat reaches noxious temperatures. After the pairing session, the neuron responded robustly to the auditory tone, and continued to do so for a median of 13 presentations of the tone (delivered every 120 seconds; Fig. 9A, range = 16-10 stimuli). After the tone responses extinguished, we delivered a second session of 15 pairings of the auditory tone and heat stimuli, after which the neuron again began responding to the auditory stimulus, and did so for the next 14 stimuli, until this auditory response extinguished (Fig. 9B, median = 13.5 stimuli, range = 20-9 stimuli). Sham pairings (see methods; “unconditioned tone”), where the auditory tone was presented without the noxious heat stimulus, had no effect on PBN activity (Fig. 9, grey traces).

We conditioned nociceptive PBN neurons to auditory stimuli in 24 neurons from 7 animals (4 males, 3 females), using the conditioning paradigm described above while measuring pupil size changes. None of these neurons responded to auditory stimuli before conditioning, but all of them began responding to the auditory stimuli after conditioning. Significant responses to auditory stimuli could be evoked for 6 to 22 stimuli (median =13) after the conditioning. All the neurons could be conditioned again after extinction of the auditory stimuli. Pupil size significantly increased after the conditioning paradigm in 10 out of the 14 neurons recorded (Fig 9C,D). These data suggest that PBN neurons that respond to noxious heat stimuli can be conditioned to respond to an innocuous auditory tone.

### Discussion Anesthesia alters properties of PBN neurons

Much of what we know about the neurophysiology of nociception and pain originates from studies using anesthetics which may alter neuronal responses to noxious stimuli. Some anesthetics nearly completely suppress responses of PBN neurons (Luo et al., 2018; Melonakos et al., 2021). We and others have found that PBN neurons do respond to noxious stimuli under urethane anesthesia (Bernard and Besson, 1990; Uddin et al., 2018; Raver et al., 2020; Uddin et al., 2021). Here we show that those responses are different from the ones recorded in awake animals. In awake animals, spontaneous PBN neuronal activity is, on average, 3 times higher and PBN responses to noxious stimuli are, on average, 5 times larger in amplitude than those in urethane anesthetized animals. In contrast, response duration was indistinguishable in awake and anesthetized animals.

### PB^CGRP^ neurons respond to noxious stimuli

PBN neurons expressing calcitonin gene-related peptide (CGRP) have been implicated in driving pain behaviors. Inhibition of PB^CGRP^ neurons attenuates pain behaviors (Han et al., 2015). PB^CGRP^ neurons are also involved in acute pain perception, as they respond to acute noxious stimuli (Campos et al., 2018; Kang et al., 2022). Our findings corroborate this conclusion, demonstrating that populations of PB^CGRP^ neurons respond robustly to thermal noxious stimuli.

One study reported that CGRP application in PBN can produce analgesia, and that this effect is blocked by sub-nanomolar concentrations of a CGRP antagonist (Wang et al., 2021). To our knowledge, this is the only report suggesting that CGRP might be anti-nociceptive. In contrast, we and others have found that PB^CGRP^ neurons robustly increase their activity in response to nociceptive stimuli.

### PBN activity correlates with internal states

Recordings from awake animals are required to determine how “internal states” might affect PBN activity and pain perception. Internal states directed towards distractions, such as cognitively demanding tasks, can affect pain perception. Detection of a noxious stimulus is compromised while performing a working memory-intensive task (Buhle and Wager, 2010). Inflammatory pain is attenuated in states of hunger, through a process involving neuropeptide Y (NPY) signaling (Alhadeff et al., 2018); manipulating NPY in PBN can mimic hunger states, suggesting a role of PBN in this state-dependent pain modulation. Similarly, suppressing PBN activity inhibits arousal (Kaur et al., 2017), whereas activating PBN neurons increases arousal (Kaur and Saper, 2019).

We found a direct correlation between PBN activity and arousal, as measured by pupil size, during presentation of a noxious stimulus. Since changes in PBN activity preceded changes in pupil size, PBN activity may be influencing internal states. A potential concern is that thermal stimulation may directly lead to pupil dilation, independent of central mechanisms. This is because thermal stimulation of the snout activates the V1 branch of the trigeminal nerve (Kim et al., 2014), potentially causing pupil dilation via the ciliary nerve, which contains fibers from the V1 branch of the trigeminal nerve (Joo et al., 2014; McDougal and Gamlin, 2015). However, our finding that pupil size and PBN activity are correlated in the absence of heat stimuli argues against this possibility.

### Nociceptive PBN neurons can be conditioned to respond to an auditory tone

In addition to its role in nociception and arousal, the PBN is also involved in aversive behaviors. For example, stimulating the PBN induces conditioned taste aversion, whereas silencing it attenuates conditioned taste aversion (Carter et al., 2015; Chen et al., 2018). Optogenetic stimulation of the PBN nucleus induces context-dependent freezing, demonstrating its involvement with threat memory and fear conditioning (Han et al., 2015; Campos et al., 2018; Bowen et al., 2020). Although these studies show that PBN neurons are necessary and sufficient for fear conditioning, they do not elucidate the underlying function of specific PBN neurons. We found that the same PBN neurons that respond to noxious stimuli can be conditioned to respond to non-noxious stimuli. Changes in pupil size mirrored this conditioned behavior. These findings suggest that conditioned aversion may involve plasticity in the response properties of PBN neurons, consistent with evidence for a causal role of the PBN in conditioned aversive behaviors (Carter et al., 2015; Campos et al., 2018; Chen et al., 2018; Bowen et al., 2020).

The auditory-nociceptive conditioning paradigm in our study evoked short-lasting conditioning of both pupil dilation responses and PBN responses to the conditioned auditory stimuli (Fig. 9). In contrast, conditioned aversion induced by pairing direct stimulation of the PBN with tastants, or pairing tastants with appetite-suppressing substances, produced aversive behaviors that can last days (Carter et al., 2015; Campos et al., 2018; Chen et al., 2018; Bowen et al., 2020). A causal role for the PBN in this long-lasting aversion is supported by finding that suppressing PBN activity attenuates the conditioned behavior (Carter et al., 2015; Campos et al., 2018; Chen et al., 2018; Bowen et al., 2020).

Our findings (Fig. 9) confirm that individual PBN neurons can respond to multiple modalities (Sammons et al., 2016; Campos et al., 2018; Bowen et al., 2020), including to nociceptive stimuli (Kang et al., 2022). Individual PBN neurons can respond during the conditioning phase, where a noxious stimulus is paired with a non-noxious tone, but they do not respond when presented with the conditioned tone 24 hours later (Kang et al., 2022). Our findings show that the same PBN neurons that respond to noxious heat can be rapidly conditioned to respond to an auditory tone, confirming that PBN neurons may be responsible for aversive memory formation. However, we also show that this conditioned response is short lived, lasting only up to one hour after conditioning, suggesting that PBN neurons may be involved in short term aversive memory retrieval but not in long term aversive memory retrieval.

## Notes

### Competing Interest Statement

The authors have declared no competing interest.

### Summary of Updates

Edited text. Removed data related to sex differences.

## References

Akintola T, Raver C, Studlack P, Uddin O, Masri R, Keller A (2017) The grimace scale reliably assesses chronic pain in a rodent model of trigeminal neuropathic pain. Neurobiol Pain, 2:13–17.

Alhadeff AL, Su Z, Hernandez E, Klima ML, Phillips SZ, Holland RA, Guo C, Hantman AW, De Jonghe BC, Betley JN (2018) A Neural Circuit for the Suppression of Pain by a Competing Need State. Cell, 173:140–152.e15.

Auvray M, Myin E, Spence C (2010) The sensory-discriminative and affective-motivational aspects of pain. Neurosci Biobehav Rev, 34:214–223.

Bennett GJ, Xie YK (1988) A peripheral mononeuropathy in rat that produces disorders of pain sensation like those seen in man. Pain, 33:87–107.

Bernard JF, Besson JM (1990) The spino(trigemino)pontoamygdaloid pathway: electrophysiological evidence for an involvement in pain processes. J Neurophysiol, 63:473–490.

Borsook D (2012) Neurological diseases and pain. Brain, 135:320–344.

Bowen AJ, Chen JY, Huang YW, Baertsch NA, Park S, Palmiter RD (2020) Dissociable control of unconditioned responses and associative fear learning by parabrachial CGRP neurons. Elife, 9:e59799.

Bruno CA, O’Brien C, Bryant S, Mejaes JI, Estrin DJ, Pizzano C, Barker DJ (2021) pMAT: An open-source software suite for the analysis of fiber photometry data. Pharmacol Biochem Behav, 201:173093.

Buhle J, Wager TD (2010) Performance-dependent inhibition of pain by an executive working memory task. Pain, 149:19–26.

Bushnell MC, Ceko M, Low LA (2013) Cognitive and emotional control of pain and its disruption in chronic pain. Nat Rev Neurosci, 14:502–511.

Cai YQ, Wang W, Paulucci-Holthauzen A, Pan ZZ (2018) Brain circuits mediating opposing effects on emotion and pain. J Neurosci, 38:6340–6349.

Campos CA, Bowen AJ, Roman CW, Palmiter RD (2018) Encoding of danger by parabrachial CGRP neurons. Nature, 555:617–622.

Carter ME, Han S, Palmiter RD (2015) Parabrachial calcitonin gene-related peptide neurons mediate conditioned taste aversion. J Neurosci, 35:4582–4586.

Castle MJ, Gershenson ZT, Giles AR, Holzbaur EL, Wolfe JH (2014) Adeno-associated virus serotypes 1, 8, and 9 share conserved mechanisms for anterograde and retrograde axonal transport. Hum Gene Ther, 25:705–720.

Chaplan SR, Bach FW, Pogrel JW, Chung JM, Yaksh TL (1994) Quantitative assessment of tactile allodynia in the rat paw. J Neurosci Methods, 53:55–63.

Chen JY, Campos CA, Jarvie BC, Palmiter RD (2018) Parabrachial CGRP Neurons Establish and Sustain Aversive Taste Memories. Neuron, 100:891–899.e5.

Chiang MC, Bowen A, Schier LA, Tupone D, Uddin O, Heinricher MM (2019) Parabrachial Complex: A Hub for Pain and Aversion. J Neurosci, 39:8225–8230.

Chiang MC, Nguyen EK, Canto-Bustos M, Papale AE, Oswald AM, Ross SE (2020) Divergent Neural Pathways Emanating from the Lateral Parabrachial Nucleus Mediate Distinct Components of the Pain Response. Neuron, 106:927–939.e5.

de Gee JW, Tsetsos K, Schwabe L, Urai AE, McCormick D, McGinley MJ, Donner TH (2020) Pupil-linked phasic arousal predicts a reduction of choice bias across species and decision domains. Elife, 9:e54014.

Deuis JR, Dvorakova LS, Vetter I (2017) Methods used to evaluate pain behaviors in rodents. Front Mol Neurosci, 10:284.

Dixon WJ (1965) The up-and-down method for small samples. Journal of the American Statistical Association, 60:967–978.

Fields HL (1999) Pain: an unpleasant topic. Pain, Suppl 6:S61–S69.

Han S, Soleiman MT, Soden ME, Zweifel LS, Palmiter RD (2015) Elucidating an Affective Pain Circuit that Creates a Threat Memory. Cell, 162:363–374.

Joo W, Yoshioka F, Funaki T, Mizokami K, Rhoton AL (2014) Microsurgical anatomy of the trigeminal nerve. Clin Anat, 27:61–88.

Kang SJ, Liu S, Ye M, Kim DI, Pao GM, Copits BA, Roberts BZ, Lee KF, Bruchas MR, Han S (2022) A central alarm system that gates multi-sensory innate threat cues to the amygdala. Cell Rep, 40:111222.

Kaur S, Saper CB (2019) Neural Circuitry Underlying Waking Up to Hypercapnia. Front Neurosci, 13:401.

Kaur S, Wang JL, Ferrari L, Thankachan S, Kroeger D, Venner A, Lazarus M, Wellman A, Arrigoni E, Fuller PM, Saper CB (2017) A Genetically Defined Circuit for Arousal from Sleep during Hypercapnia. Neuron, 96:1153–1167.e5.

Kim YS, Chu Y, Han L, Li M, Li Z, Lavinka PC, Sun S, Tang Z, Park K, Caterina MJ, Ren K, Dubner R, Wei F, Dong X (2014) Central terminal sensitization of trpv1 by descending serotonergic facilitation modulates chronic pain. Neuron,

Langford DJ, Bailey AL, Chanda ML, Clarke SE, Drummond TE, Echols S, Glick S, Ingrao J, Klassen-Ross T, Lacroix-Fralish ML, Matsumiya L, Sorge RE, Sotocinal SG, Tabaka JM, Wong D, van den Maagdenberg AM, Ferrari MD, Craig KD, Mogil JS (2010) Coding of facial expressions of pain in the laboratory mouse. Nat Methods, 7:447–449.

Legrain V, Crombez G, Mouraux A (2011) Controlling attention to nociceptive stimuli with working memory. PLoS One, 6:e20926.

Luo T, Yu S, Cai S, Zhang Y, Jiao Y, Yu T, Yu W (2018) Parabrachial Neurons Promote Behavior and Electroencephalographic Arousal From General Anesthesia. Front Mol Neurosci, 11:420.

Matsumoto N, Bester H, Menendez L, Besson JM, Bernard JF (1996) Changes in the responsiveness of parabrachial neurons in the arthritic rat: an electrophysiological study. J Neurophysiol, 76:4113–4126.

McDougal DH, Gamlin PD (2015) Autonomic control of the eye. Compr Physiol, 5:439–473.

Melonakos ED, Siegmann MJ, Rey C, O’Brien C, Nikolaeva KK, Solt K, Nehs CJ (2021) Excitation of Putative Glutamatergic Neurons in the Rat Parabrachial Nucleus Region Reduces Delta Power during Dexmedetomidine but not Ketamine Anesthesia. Anesthesiology, 135:633–648.

Melzack R, Casey KL (1968) Sensory, motivational and central control determinants of pain. In: The Skin Senses (Kenshalo DR, ed), pp 423–439. Springfield: Thomas.

Okubo M, Castro A, Guo W, Zou S, Ren K, Wei F, Keller A, Dubner R (2013) Transition to persistent orofacial pain after nerve injury involves supraspinal serotonin mechanisms. J Neurosci, 33:5152–5161.

Palmiter RD (2018) The Parabrachial Nucleus: CGRP Neurons Function as a General Alarm. Trends Neurosci, 41:280–293.

Price DD (2000) Psychological and neural mechanisms of the affective dimension of pain. Science, 288:1769–1772.

Raver C, Uddin O, Ji Y, Li Y, Cramer N, Jenne C, Morales M, Masri R, Keller A (2020) An Amygdalo-Parabrachial Pathway Regulates Pain Perception and Chronic Pain. J Neurosci, 40:3424–3442.

Sammons JD, Weiss MS, Victor JD, Di Lorenzo PM (2016) Taste coding of complex naturalistic taste stimuli and traditional taste stimuli in the parabrachial pons of the awake, freely licking rat. J Neurophysiol, 116:171–182.

Smith JG, Elias LA, Yilmaz Z, Barker S, Shah K, Shah S, Renton T (2013) The psychosocial and affective burden of posttraumatic neuropathy following injuries to the trigeminal nerve. J Orofac Pain, 27:293–303.

Stringer C, Pachitariu M, Steinmetz N, Reddy CB, Carandini M, Harris KD (2019) Spontaneous behaviors drive multidimensional, brainwide activity. Science, 364:255.

Treede RD, Apkarian AV, Bromm B, Greenspan JD, Lenz FA (2000) Cortical representation of pain: functional characterization of nociceptive areas near the lateral sulcus. Pain, 87:113–119.

Turnes JM, Araya EI, Barroso AR, Baggio DF, Koren LO, Zanoveli JM, Chichorro JG (2022) Blockade of kappa opioid receptors reduces mechanical hyperalgesia and anxiety-like behavior in a rat model of trigeminal neuropathic pain. Behav Brain Res, 417:113595.

Uddin O, Anderson M, Smith J, Masri R, Keller A (2021) Parabrachial complex processes dura inputs through a direct trigeminal ganglion-to-parabrachial connection. Neurobiol Pain, 9:100060.

Uddin O, Studlack P, Akintola T, Raver C, Castro A, Masri R, Keller A (2018) Amplified parabrachial nucleus activity in a rat model of trigeminal neuropathic pain. Neurobiol Pain, 3:22–30.

Wang LL, Wang HB, Fu FH, Yu LC (2021) Role of calcitonin gene-related peptide in pain regulation in the parabrachial nucleus of naive rats and rats with neuropathic pain. Toxicol Appl Pharmacol, 414:115428.

Wu TH, Hu LY, Lu T, Chen PM, Chen HJ, Shen CC, Wen CH (2015) Risk of psychiatric disorders following trigeminal neuralgia: a nationwide population-based retrospective cohort study. J Headache Pain, 16:64.

Yeomans DC, Proudfit HK (1996) Nociceptive responses to high and low rates of noxious cutaneous heating are mediated by different nociceptors in the rat: electrophysiological evidence. Pain, 68:141–150.

